# Trait matching without traits: using correspondence analysis to investigate the latent structure of interaction networks

**DOI:** 10.1101/2024.10.22.619454

**Authors:** Lisa Nicvert, Hervé Fritz, Stéphane Dray

## Abstract

Species interactions in ecological communities are often represented as networks, the structure of which is thought to be linked to species’ interaction niches (or Eltonian niches). Interaction niches are intimately related to the notion of trait matching, which posits that a species interacts preferentially with partners whose traits are complementary to their own.

Multivariate methods are commonly used to quantify species environmental niches (or Grinnellian niches). More recently, some of these methods have also been used to study the interaction niche, but they consider only the niche optimum and require trait data.

In this article, we use the correspondence analysis (CA) framework to study interaction networks and investigate trait matching without requiring trait data, using the notion of latent traits. We use reciprocal scaling, a method related to CA, to estimate niche optima and breadths, defined respectively as the mean and standard deviation of the latent traits of species’ interacting partners. We present the method, test its performance using a simulation model we designed, and analyze a real frugivory network between birds and plants.

The simulation study shows that the method is able to recover niche breadths and optima for data generated with parameters typical of ecological networks. The birds-plants network analysis shows strong relationships between species latent traits and niche breadths: a posteriori correlation with measured traits suggests that birds and plants of intermediate size tend to have the broadest niches. Additionally, birds preferentially foraging in the understory have broader niches than birds preferentially foraging in the canopy.

CA and reciprocal scaling are described as fruitful exploratory methods to characterize species interaction profiles, provide an ecologically meaningful graphical representation of interaction niches, and explore the effect of latent traits on network structure.

## Introduction

The ecological niche constitutes the pool of environmental conditions and resources required for the persistence of species (Hutchinson, 1957). Species’ abiotic requirements are described with the Grinnellian niche (Grinnell, 1924), which centers on species’ physical and environmental needs. In contrast, species’ biotic requirements are described with the Eltonian niche (originally formalized in the context of food webs by Elton, 1927), which sees species’ niche through the lens of their interactions with other organisms. Because it encapsulates both abiotic and biotic requirements, the niche concept underpins our understanding of species’ distributions, coexistence and competitive exclusion (Chase and Leibold, 2003).

The niche is often characterized by its optimum and breadth. Niche optimum describes the conditions under which species growth is maximized (Kermavnar et al., 2023; Treurnicht et al., 2020), whereas niche breadth defines the range of conditions tolerated by a species (Carscadden et al., 2020; Sexton et al., 2017). Species with a broad niche are called generalists, and those with a narrow niche are called specialists. A common approach to describe species’ niche space is to use Gaussian curves along each axis (Gauch Jr. and Whittaker, 1972). In this framework, the mean represents the niche optimum, and the standard deviation the niche breadth, along each dimension.

The methods used to describe and analyze niches are often different depending on whether the focus is on the environmental or on the interaction niche. Interaction niches are usually analyzed through the formalism of ecological networks of species (nodes) and interactions (edges) (Bascompte and Jordano, 2007; Ings et al., 2009). In these networks, niche breadth has most often been defined as the number of interacting partners, potentially considering their abundance (Devictor et al., 2010). Alternatively, it has also been defined as the range of traits of interacting partners (Dehling, Jordano, et al., 2016; Dehling, Töpfer, et al., 2014; Maglianesi et al., 2015).

Environmental niches are commonly estimated with multivariate ordination methods, which allow to estimate niche optima along environmental gradient(s). The basis of these methods is weighted averaging (Whittaker, 1956): it consists in averaging the values of the environmental variable over the samples in which a species occurs, weighted by the species’ abundance, to estimate its niche optimum. These multivariate methods can use implicit or explicit environmental gradients, and optionally relate these gradients with species traits. When the only data available are species abundances sampled in different sites, correspondence analysis (CA) (Hill, 1973, 1974), estimates species’ niche optima on latent gradients. When environmental variables are recorded, canonical correspondence analysis (ter Braak, 1986) estimates species niche optima on environmental gradients. Finally, when traits and environmental variables are available, one can use fourth-corner analysis (Legendre, Galzin, et al., 1997), RLQ (Dolédec et al., 1996) and double constrained correspondence analysis (ter Braak et al., 2018) to link niche optima estimated on environmental gradients to species traits.

In this study, we focus on characterizing species interactions in ecological networks. We also focus on realized niches (that is, the portion of niche space species effectively occupy) estimated from community data. It is generally accepted that ecological interactions do not arise randomly, but reflect multiple drivers such as neutral effects, trait matching, and evolutionary history (Peralta et al., 2024; Vázquez et al., 2009). Neutral effects imply that a given species is more likely to interact with abundant than with rare species (Vázquez et al., 2009). Trait matching posits that species with complementary traits preferentially engage in interactions. For instance, body size has been identified as a key factor influencing food web structure (Elton, 1927; Ings et al., 2009). Species evolutionary history is also thought to influence their interaction patterns, for instance by constraining traits via phylogenetic inertia or coevolution (Benadi et al., 2022; Dormann et al., 2017). Finally, these processes are not mutually exclusive and often interact to shape observed patterns (Vázquez et al., 2009).

Recently, several multivariate methods have been applied to study interaction networks. In particular, many methods have investigated trait matching: for instance, RLQ and fourth-corner analyses have been used to measure the correlation between two sets of partner species’ traits, weighted by the interaction matrix (Albrecht et al., 2018; Bender et al., 2018; Dehling, Töpfer, et al., 2014).

Classical methods to analyze ecological networks have brought invaluable insights on trait matching and degree of specialization, but also have shortcomings. First, trait matching in ecological networks has mainly been investigated separately from interaction niches (Godoy et al., 2018; Phillips et al., 2020). However, two species only likely interact if their niches match (e.g. in the simulations models of Benadi et al., 2022; Fründ et al., 2016), so that trait matching implicitly relies on species interaction niche (Albrecht et al., 2018; Eklöf et al., 2013). Second, trait matching studies only consider the average relationship between species traits across the network. Hence, they generally ignore the variability around trait matching (i.e., niche breadth). Lastly, methods to study trait matching require the collection of trait data, which has been recognized as a challenge (Vázquez et al., 2009), as it is tedious to collect multiple traits and identify a priori which ones are relevant drivers of interactions.

Here, we propose correspondence analysis (CA) (Hill, 1974; Hirschfeld, 1935), a method often used to study species’ environmental niches, to study species’ interaction niches in ecological networks. We interpret ordination scores produced by CA as species’ latent traits, corresponding to the optima of their interaction niches. We also use reciprocal scaling (Thioulouse and Chessel, 1992) to estimate interaction niches breadths, defined as the standard deviation of the latent traits of a species’ interacting partners. Our framework also allows to measure the magnitude and dimensionality of trait matching, without using trait data. To do so, we use the notion of latent traits, which are understood as a proxy of various unmeasured species characteristics. Lastly, these methods also enable ecologically meaningful graphical representation of species interaction niches.

In this article, we first present CA and reciprocal scaling and their interpretation in the context of interaction networks. Then, we evaluate the performance of these methods to estimate species niches (optimum and breadth) using simulated data. Finally, we analyze a real birds-plants interaction network (Dehling, Bender, et al., 2021) to show how our framework can help measure and represent interaction niches, highlight trait matching, and explore the drivers of niche breadth.

## Material and methods

All analyses were performed with R 4.4.2 (R Core Team, 2024) and are available at https://doi.org/10.6084/m9.figshare.28625270.v3 (Nicvert et al., 2025).

### Notations

In this part, we consider a *r × c* matrix **Y** = [*y*_*ij*_] representing the bipartite interaction network between *r* resource species and *c* consumer species (either abundances or presences). We use the terms “resource” and “consumer” in accordance with the literature, as these terms can encompass diverse bipartite network types (Fründ et al., 2016; Ings et al., 2009).

For the following mathematical developments, we define **P** as the table of relative frequencies of interactions: **P** = [*y*_*ij*_*/y*_++_] (where 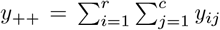 is the grand total of **Y**). We also define the weight matrices **D**_**r**_ = diag(**r**) for resources (rows) and **D**_**c**_ = diag(**c**) for consumers (columns), where the vectors **r** = **P1**_**r**_ = [*p*_1+_, …, *p*_*r*+_]^*⊤*^ and **c** = **P**^*⊤*^**1**_**c**_ = [*p*_+1_, …, *p*_+*c*_]^*⊤*^ represent, respectively, the rows and columns marginal sums (i.e. 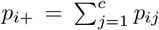 and 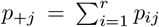). Throughout this article, ^*⊤*^ will denote matrix transposition.

### Quantify trait matching

We consider two known traits **x** and **y** measured, respectively, on the resource and consumer species. A simple measure of trait matching is given by the fourth-corner statistic cor_**P**_(**x, y**) (Legendre, Galzin, et al., 1997). It measures the correlation between the traits of consumers and resources, weighted by their interaction frequencies:

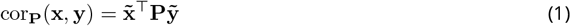

where 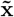 and 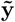 are centered scaled versions of the traits, respectively using weights **D**_**r**_ and **D**_**c**_. If centered traits **y**_**0**_ and **y**_**0**_ are considered instead, the product **x**_**0**_^*⊤*^**Py**_**0**_ represents a covariance denoted cov_**P**_(**x, y**).

This fourth-corner statistic has been used to quantify trait matching in interaction networks (Albrecht et al., 2018; Bender et al., 2018). However, when the traits contributing to the matching are not known, it cannot be computed. In that case, indirect gradient methods, like correspondence analysis, can be useful to uncover latent axes that structure interactions in ecological networks.

### Correspondence analysis (CA) for interaction networks

Correspondence analysis (Hill, 1974; Hirschfeld, 1935) is a multivariate method used to analyze contingency tables. In the following, we describe CA applied to contingency tables of interaction counts.

CA is based on the generalized singular value decomposition of the matrix **D**_**r**_^*−*1^**P**_**0**_**D**_**c**_^*−*1^, where **P**_**0**_ = **P** − **rc**^*⊤*^ is the double centered matrix of interaction frequencies **P**:

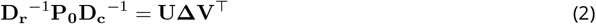

The matrices **U** and **V** are orthonormal with respect to weights **D**_**r**_ and **D**_**c**_ (**U**^*⊤*^**D**_**r**_**U** = **I** and **V**^*⊤*^**D**_**c**_**V** = **I**) and contain respectively the left (resource) and right (consumer) generalized singular vectors. **Δ** is the diagonal matrix containing ordered singular values. The matrix of CA eigenvalues **Λ** is equal to **Δ**^2^.

To interpret these results, let us consider a given dimension *k*. The previous equation can then be rewritten as:

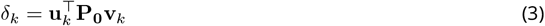

As the matrix **P**_**0**_ is double centered (by rows and columns), its singular vectors stored in **U** and **V** are centered as well. It follows that 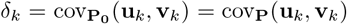. Moreover, singular vectors have also unit variance (orthonormality constraint). Hence, Equation (3) is a correlation and 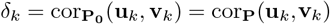. This is strictly analogous to the trait matching equation (1), but known traits 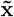 and 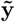 are replaced with the generalized singular vectors **u**_*k*_ and **v**_*k*_ in Equation (3). Therefore, CA amounts to finding the singular vectors **u**_*k*_ and **v**_*k*_ that are maximally correlated. Consequently, the scores **u**_*k*_ and **v**_*k*_ can be interpreted as latent traits, associated to resources (respectively, consumers), which maximize matching between these latent traits.

Using the relationship between **Δ** and **Λ**, we can show that the square root of a CA eigenvalue 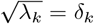 represents the absolute value of the correlation between **u**_*k*_ and **v**_*k*_: an eigenvalue close to one indicates that latent traits can correctly explain the probability of interaction. The global relationship between all pairs of latent traits can also be tested statistically, using a *χ*^2^ test on the interaction matrix (Abdi and Béra, 2017; Beh, 2004). This test can be performed analytically if its assumptions are met, but in the context of interactions networks, a permutation approach will often be better suited.

CA can assess trait matching, without using trait data, using these latent scores. However, these latent traits represent more than species unmeasured traits. Indeed, they can be interpreted in a broader sense as proxies for unmeasured properties for species, beyond their traits (for instance, phylogenetic signal like in Benadi et al., 2022). To facilitate interpretation, it is possible to link latent scores identified by CA to known properties, such as traits or phylogeny, a posteriori.

CA assigns similar scores to species with the same interaction profile: in other words, two species interacting with the same partners will be positioned nearby in the CA multivariate space. Species scores can therefore be used to reorder rows and columns of the interaction matrix and highlight the structure of the network (Lewinsohn et al., 2006). Appendix B shows an example of reordering with the birds-plants interaction network used in this article.

Standard outputs of CA consist of two biplots in which both consumers and resources are displayed in the same multivariate space. To convert consumer scores in the resource space (and vice versa), we use the following transition formulas:

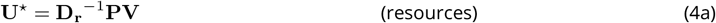

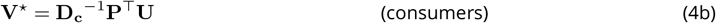

Equation (4a) expresses resources’ scores as a weighted mean of the scores of consumers they interact with. Reciprocally, Equation (4b) expresses consumers’ scores as a weighted mean of the scores of resources they interact with. The transformation described in Equations (4) is called weighted averaging.

The scores **U**^*⋆*^ and **V**^*⋆*^ can also be computed using scores **U** (respectively, **V**) scaled by the square roots of eigenvalues (Legendre and Legendre, 2012):

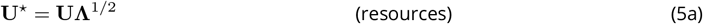

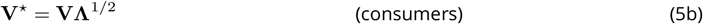

**U**^*⋆*^ and **V**^*⋆*^ can be interpreted as niche optima, defined as the mean value of a species’ interacting partners’ latent traits (Equation (4)). Equivalently, **U**^*⋆*^ and **V**^*⋆*^ can be interpreted as species’ latent traits, as they are proportional to **U** and **V** (Equation (5)). To sum up, the latent traits of resources and consumers estimated with CA are mathematically related, as they are estimated by maximizing their matching.

Resources and consumers can be plotted in two spaces, corresponding to two biplots. A first biplot can be drawn by displaying resources’ scores given by their latent trait (**U**) and consumers’ scores given by weighted averaging (**V**^*⋆*^). The second biplot mirrors the first, with consumers’ scores **V** (latent traits) and resources’ scores **U**^*⋆*^ (weighted averaging).

These scores make CA particularly suitable for analyzing interaction networks. However, this method summarizes species’ niches with their optima, and completely ignores niche breadth. Moreover, the niche optima can only be visualized on two different biplots.

### Reciprocal scaling

Reciprocal scaling, proposed by Thioulouse and Chessel (1992), is an extension of CA that solves the two issues mentioned above. First, this method provides niche breadths, in addition to optima. Second, it defines a common multivariate space for resource and consumers. Reciprocal scaling shifts the focus of the analysis by considering the interactions, rather than the species, as statistical individuals. For that, it uses the interpretation of CA as a special case of canonical correlation analysis (for complete mathematical development, see Thioulouse and Chessel, 1992).

Reciprocal scaling defines a score *h*_*k*_(*i, j*) for each interaction between species *i* and *j* in the *k*-th dimension of the multivariate space (note that it is defined only if species *i* and *j* interact). These interactions correspond to non-empty cells in the interaction matrix **Y** (they are called correspondence in Thioulouse and Chessel, 1992). Interactions scores can be computed from CA scores using the following formula:

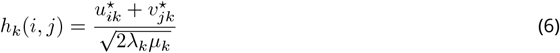

where *λ*_*k*_ is the CA eigenvalue for axis *k* and 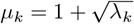. The scores *h*_*k*_(*i, j*) can be grouped into a vector **h**_*k*_ of length 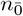 for dimension *k* (where 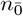 is the number of unique interactions in the matrix, i.e. non-empty cells).

Using the interactions scores *h*_*k*_(*i, j*), it is possible to display the interaction niche of a species as a cloud of interactions in the multivariate space. The niche can then be summarized by estimating its optimum and breadth using a Gaussian approximation. If we consider species *s* in the multivariate plane given by axes *k* and *l*, its niche is approximated by an ellipse (see Figure 1). The center of the ellipse represents the niche optima of species *s* on axes *k* and *l* (weighted averages of interaction scores). The semi-minor and -major axes represent its niche breadths on axes *k* and *l* (weighted standard deviations). Finally, the orientation of the ellipse represents the weighted covariance for axes *k* and *l*.

**Figure 1.**
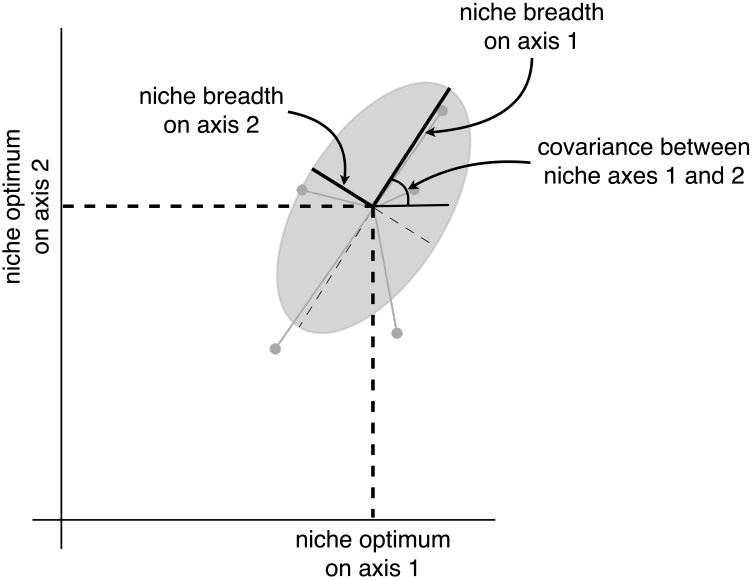
Visualization of the interaction niche in a 2-dimensional space. The niche is depicted by an ellipse whose center represents niche optima and semi-axes represent niche breadths. The angle of the ellipse represents the covariance between niche axes. Points represent interaction scores.

Formulas to compute niche optima, breadths and covariances are given in Appendix C (from Thioulouse and Chessel (1992), and adapted with the notation used in this article). We can show that the niche optima computed from these scores correspond to the scores **U**^*⋆*^ and **V**^*⋆*^ defined above, up to a scaling factor (see Equations (C.1) and (C.2) in Appendix C).

### Simulation

We simulated interaction data to evaluate the performance of CA and reciprocal scaling in estimating parameters of species interaction niches. Below, we describe the simulation model and the different experiments we conducted.

#### Simulation model

Our simulation model takes into account species traits (through trait matching) and their abundances (through neutral effects) and combines these two processes to generate an observed interaction matrix (see Figure 2). This model is inspired by the models of Fründ et al. (2016) and Benadi et al. (2022) for the simulation of trait-based interaction networks, and by the models of Dray and Legendre (2008) and Minchin (1987) for niche modeling in the context of species-sites associations.

**Figure 2.**
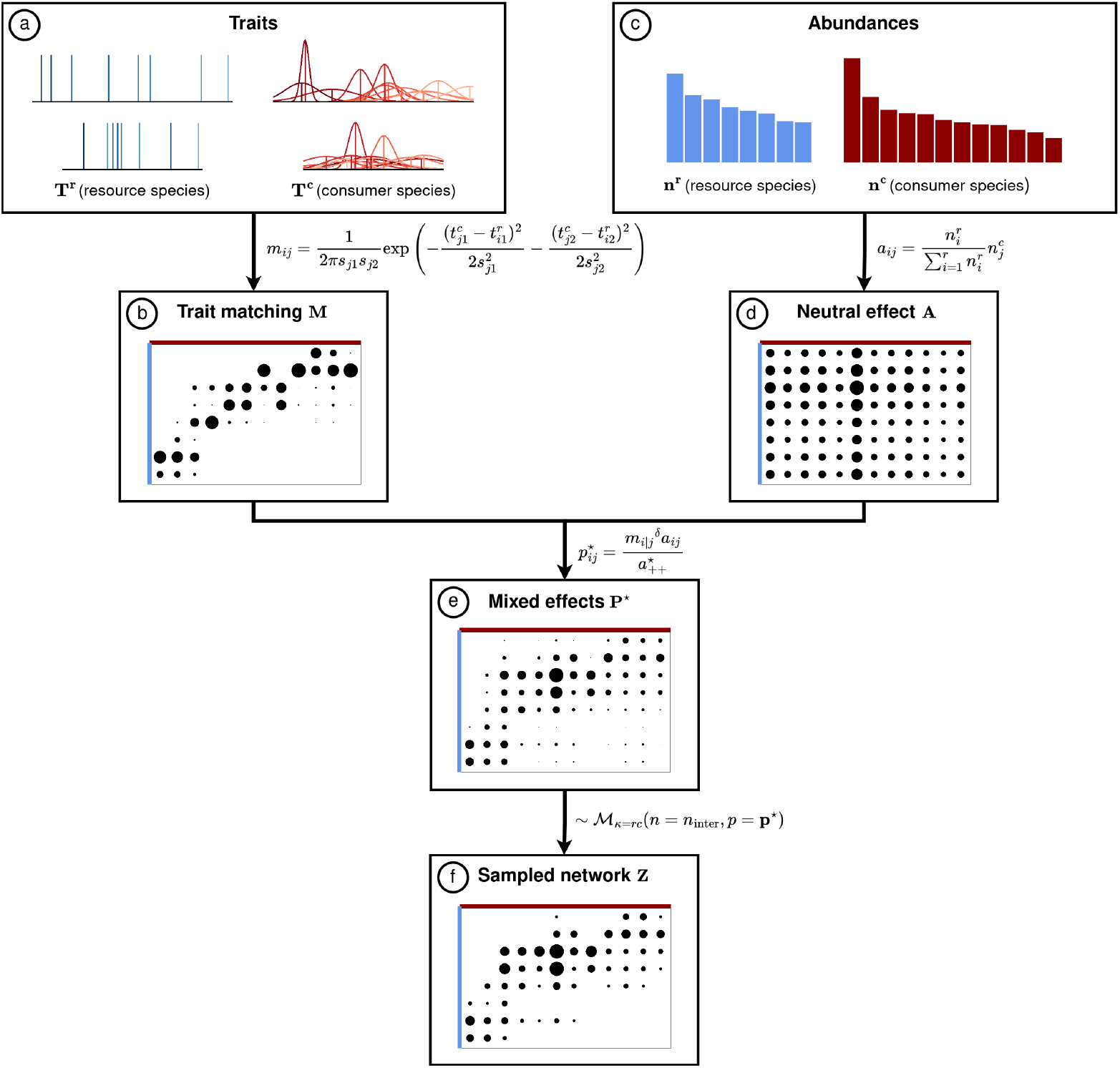
Model used to simulate interactions between consumer and resource species. (a) Two traits are generated for each species, corresponding to their niche optima. For consumers, standard deviations are also associated to traits to generate the niche breadths. (b) Interaction probabilities due to trait matching are computed from species traits. (c) In parallel, abundances of resources and consumers are generated. (d) Interactions counts representing the neutral effect of abundances on interaction probability are computed from the abundances. (e) Matching and abundance-driven interactions are combined to get the interactions counts based on both processes. (f) Observed interactions are then sampled from the mixed interaction probabilities. Equations are explained in the main text.

We consider the interactions between the *i*-th resource (*i* = 1 … *r*) and the *j*-th consumer species (*j* = 1 … *c*). The simulation procedure consists of different steps described below and illustrated in Figure 2:

**Step (a)**: we simulate two traits for both consumer and resource species (Figure 2a). The traits of resource species are stored in matrix 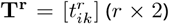 and the traits of consumers are stored in matrix 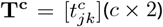. The first trait for both resources and consumers is drawn from a uniform distribution between 0 and The second trait is generated similarly, but in the interval 15-85 (see length of the traits axes in Figure 2a), so that the second trait has less weight in trait matching. For consumers, we define niche breadth with the values of **S** = [*s*_*jk*_] (shown in Figure 2a as the standard deviation of a Gaussian curve). The elements of **S** contain the absolute values of numbers drawn from a normal distribution with mean *µ*_breadth_ and standard deviation *σ*_breadth_.

**Step (b)**: we compute an interaction probability due to the matching of traits generated in the previous step (Figure 2b). This probability *m*_*ij*_ follows a bivariate normal distribution. Its mean vector is the difference between the trait values of resource species *i* and consumer species 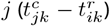; variances in the variance-covariance matrix correspond to the degree of generalization of consumer species 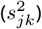, and covariances are null:

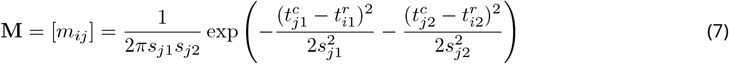

**Step (c)**: we generate species abundances for consumer and resource species (Figure 2c) from a uniform or from a log-normal distribution (see Table 1 for more details). These abundances are stored in vectors **n**^**c**^ (*c ×* 1) for consumers and **n**^**r**^ (*r ×* 1) for resources.

**Table 1.**
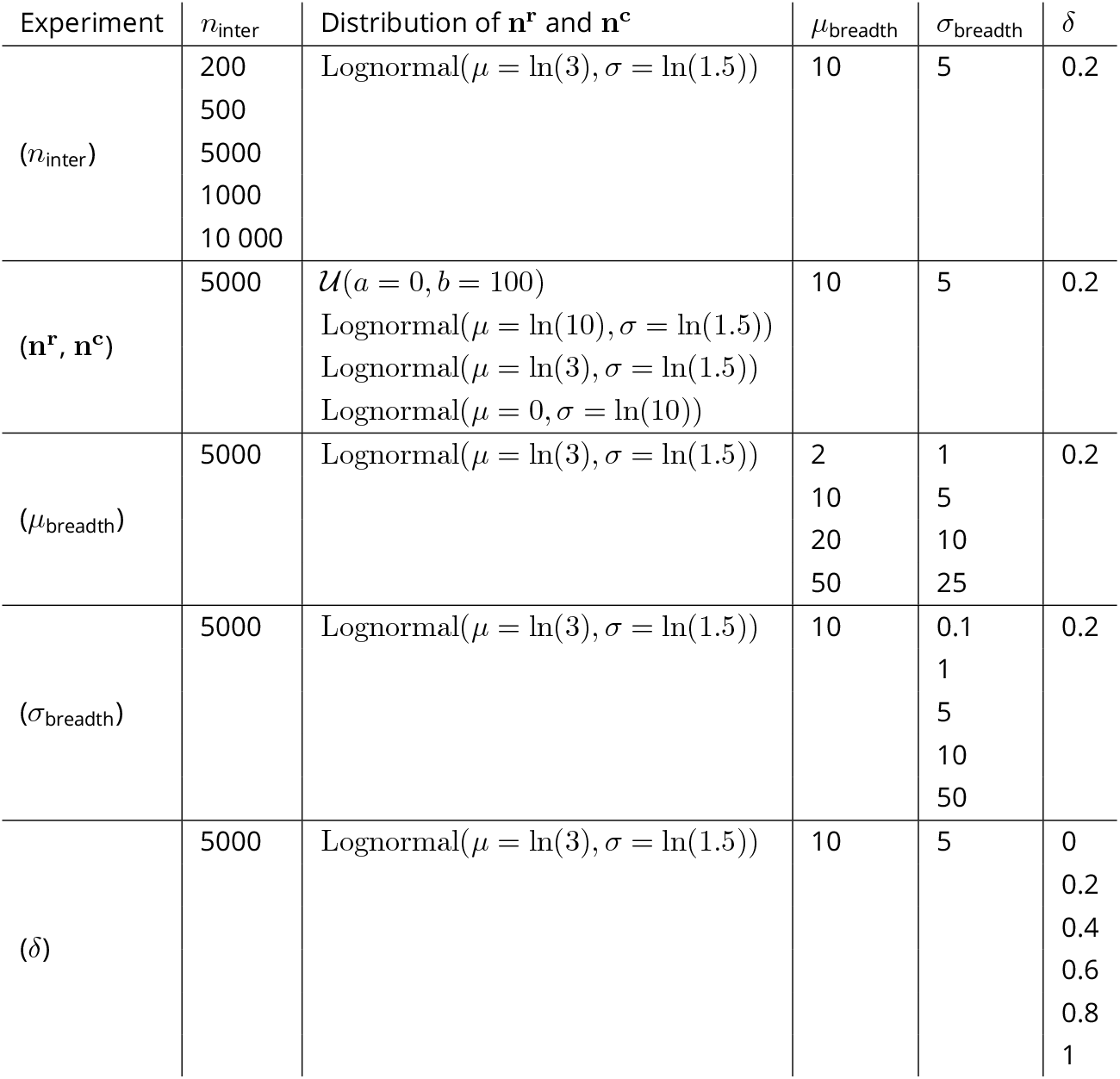
Parameter values used for the simulation study. Experiment (*n*_inter_) varies the total number of interactions. Experiment (**n**^**r**^,**n**^**c**^) progressively increases the skewness of the species abundances. Experiment (*µ*_breadth_) varies the mean value of the consumers’ niche breadth. Experiment (*σ*_breadth_) varies the standard deviation of the consumers’ niche breadth. Experiment (*δ*) varies the strength of trait matching.

**Step (d)**: we then compute a matrix of predicted interaction counts based solely on abundances (Figure 2c):

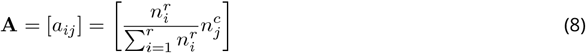

Here, we model interactions as the result of consumer choices, driven only by the relative abundance of resources (representing the availability of resource species).

**Step (e)**: we compute a composite interaction probability resulting from the combined effects of trait matching (**M**, equation (7)) and neutral abundance effects (**A**, equation (8)) (Figure 2e). A factor *δ* (0 *≤ δ ≤* 1) allows to control the relative weight of the matching and the abundance processes. If *δ* = 0, then the strength of the matching is null, and interactions are driven only by abundances. If *δ* = 1, the strength of matching is maximal, and abundances and matching combine to produce the observed pattern. The mixed interaction frequency matrix 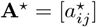 is computed as:

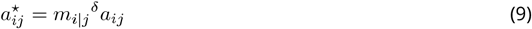

where *m*_*i*|*j*_ = *m*_*ij*_*/m*_+*j*_ represents the probability of interaction with resources per consumer species and 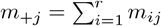 is the marginal probability of interaction for species *j*.

The interaction probability matrix 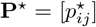 is then computed from the mixed interaction frequency matrix:

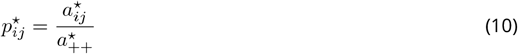

where 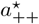 is the grand total of **A**^*⋆*^.

**Step (f)**: finally, we sample observed interactions counts from the interaction probability matrix **P**^*\**^ (Figure 2f). To do so, we sample *n*_inter_ interactions from a multinomial distribution with *κ* = *rc* outcomes corresponding to pairwise interactions. The probability vector **p**^*⋆*^ (of length *κ*) corresponds to the flattened matrix **P**^*⋆*^.

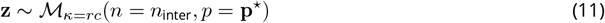

The interactions counts vector **z** is finally rearranged in the matrix **Z** (*r* × *c*) which corresponds to the sampled interaction network.

#### Simulation parameters

To evaluate the performance of our method in estimating niche parameters, we conducted 5 experiments using the model described above (detailed parameter values are presented in Table 1):

1. Experiment (*n*_inter_): we vary the total number of interactions in the matrix *n*_inter_, which represents the sampling intensity. This allows to evaluate the robustness to incomplete sampling.
2. Experiment (**n**^**r**^, **n**^**c**^): we sample **n**^**r**^ and **n**^**c**^ from a uniform distribution or from log-normal distributions either mildly, moderately or very skewed. The aim is to evaluate the robustness of the method to skewed abundances typically encountered in ecological communities.
3. Experiment (*µ*_breadth_): we vary the mean of the normal distribution in which the standard deviations of consumers’ traits *s*_*jk*_ (*k* = 1, 2) are sampled. This allows to contrast communities of generalist (high standard deviation) and specialized consumers (low standard deviation).
4. Experiment (*σ*_breadth_): we vary the standard deviation of the normal distribution in which the standard deviations of consumers’ traits *s*_*jk*_ (*k* = 1, 2) are sampled. This allows to contrast communities with homogeneous consumers (same degree of specialization) to communities with heterogeneous consumers (diverse degrees of specialization).
5. Experiment (*δ*): we vary the value of the parameter *δ*, which determines the strength of trait matching compared to neutral interactions, driven by species abundances (Equation (9)).

Each experiment described above varies one parameter of interest, and other parameters are fixed to a default value (see Table 1). In experiment (*µ*_breadth_), the standard deviation is also changed to keep a constant ratio of *σ*_breadth_ = *µ*_breadth_*/*2. For all simulations, we generated interactions between *r* = 50 resources and *c* = 60 consumers.

For each experiment, we generated 100 datasets. For each dataset, we computed the true niche optimum as the mean of the traits of the interacting partners of a species, weighted by their interaction frequencies. Similarly, the true niche breadth is computed as the weighted standard deviation of the traits of species’ interacting partners. These true values quantify species’ realized niche (inference of the fundamental niche is discussed in Appendix A). Then, we performed reciprocal scaling on each dataset using the R package ade4 (Thioulouse, Dray, et al., 2018), thus estimating the realized niches optima and breadths. To evaluate the method’s performance, we measured the correlation between true niche parameters and reciprocal scaling estimates.

### Real data analysis

#### Dataset

To illustrate CA and reciprocal scaling on real data, we analyzed a birds-plants interaction network from the ANDEAN frugivory dataset (Dehling, Bender, et al., 2021, network Peru1 from the dataset). In this dataset, consumers are birds, and resources are the fruiting plants they feed on and whose seeds they disperse. This network was sampled in the lower montane rainforest in Peru. Data were collected by direct repeated observation conducted throughout an entire year (4 times between 2009 and 2010) in 3 plots of 100 m × 30 m. A transect was used to determine focus fruiting plants inside each plot, and seed removal by birds was recorded on these plants.

In addition to interactions, this dataset also includes some species traits thought to play an important role in trait matching. To interpret the latent axes of our analysis, we used three plant traits: fruit diameter, crop mass (mean number of fruits per plant multiplied by mean fruit mass) and plant height. For birds, we also used three traits: bill width, body mass and Kipp’s index. The Kipp’s index is the Kipp’s distance, i.e. the distance from the tip of the first secondary to the wing tip, divided by wing length (Dehling, Töpfer, et al., 2014). A low Kipp’s index indicates rounded wings, and a high Kipp’s index indicates pointed wings. Plant traits were collected in the field, and bird traits on museum specimens.

Before data analysis, we filtered out the birds or plants that interact only with one other species (8 birds and 12 plants). Indeed, since CA is based on species ordination, these species are problematic from a methodological point of view (Greenacre, 2013). These very specialized species can be seen as interaction modules, which should be analyzed separately when using CA (van Dam et al., 2021). The final network has 40 plants and 53 birds (species names and traits values are listed in Appendix D).

#### Analyses

We performed CA and reciprocal scaling on this dataset, using the R package ade4 (Thioulouse, Dray, et al., 2018). We tested the relationship between latent traits using a *χ*^2^ test, and evaluated the effect size using Cramér’s *V* (Ellis, 2010).

To show how our method can answer ecological questions, we modeled the relationship between niche breadths and latent traits for resource and consumers, on 2 multivariate axes each (4 models in total). To keep simple models while also allowing a flexible shape, we considered linear and quadratic regression models. We chose the best model with a likelihood ratio test.

Finally, to interpret CA and reciprocal scaling more easily, we computed a posteriori correlations between the measured species traits and the first two latent traits.

## Results

### Simulation

The results of the 5 experiments testing the performance of niche inference are presented in Figure 4 for trait matching strength (*δ*), and Figure 3 for other parameters. First, as expected, performance improves consistently with sampling completeness for all niche measures (Experiment (*n*_inter_), Figure 3a). Second, when there are enough data (here, more than 5000 sampled interactions), the niche optima are recovered correctly on both axes (median correlation value above 0.875). Niche breadths are well-recovered for consumers (median above 0.75), but less precisely for resources (median above 0.25 only) (Figure 3a). We can also note that for all experiments, niche parameters are always better recovered on the first axis, which is consistent with the fact that, by construction, trait matching with the first trait explains more structure.

**Figure 3.**
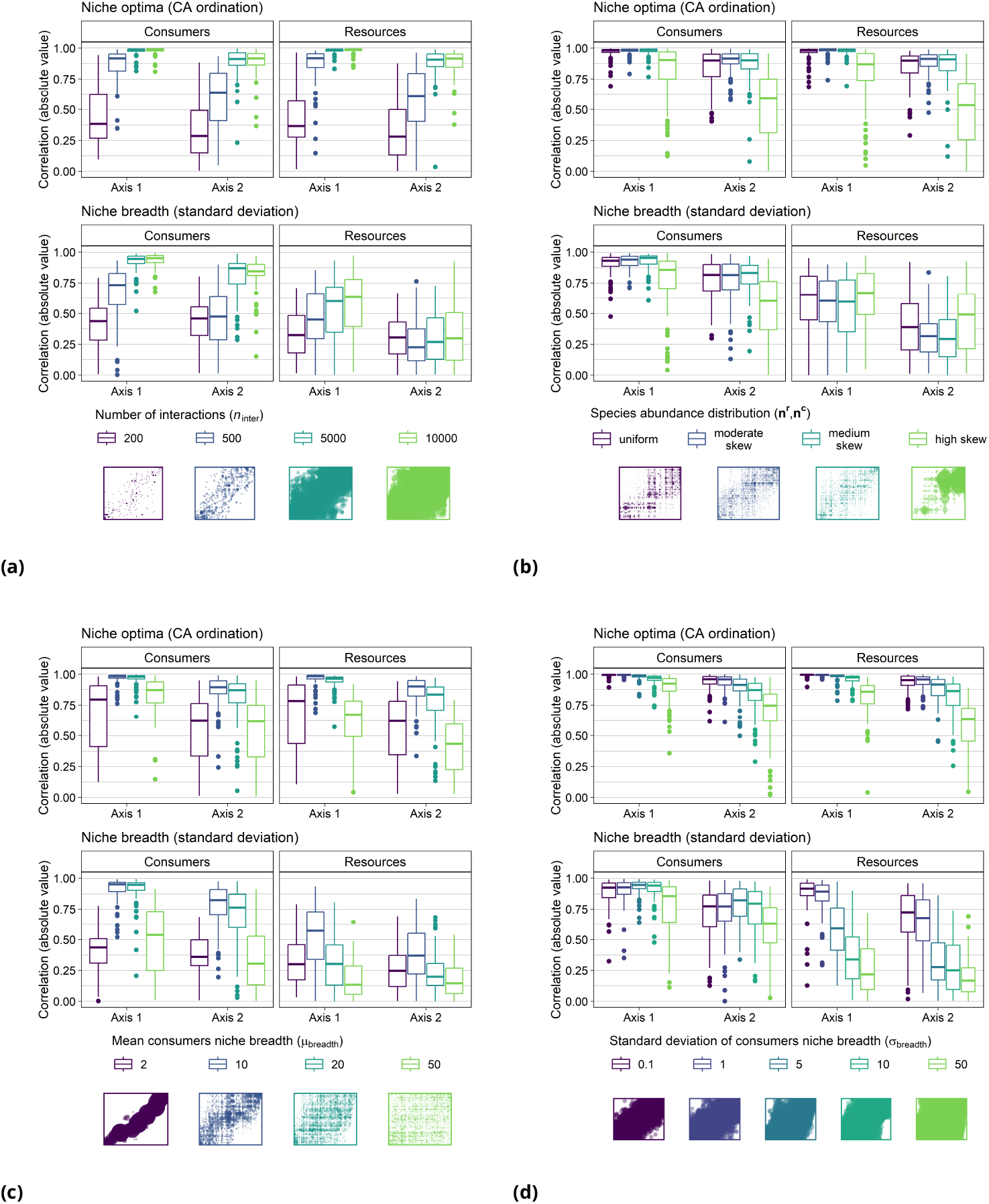
Results of the simulation study. Each subfigure explores the influence of one parameter on the model performance. (a) Effect of sampling intensity (number of interactions) (*n*_inter_). (b) Effect of species abundance distributions (**n**^**r**^, **n**^**c**^). (c) Effect of the mean consumers niche breadths (*µ*_breadth_). (d) Effect of heterogeneity (standard deviation) of consumers’ niche breadths (*σ*_breadth_). The y-axis is the absolute value of the correlation between true and estimated values of niche optima and niche breadths (respectively, top and bottom of each subplot). Below the legend, matrices exemplify network data (resources in rows, consumers in columns) generated with the corresponding parameter value. Points represent non-null interactions, with size proportional to the number of observed interactions censored at the 1st and 99th percentiles.

**Figure 4.**
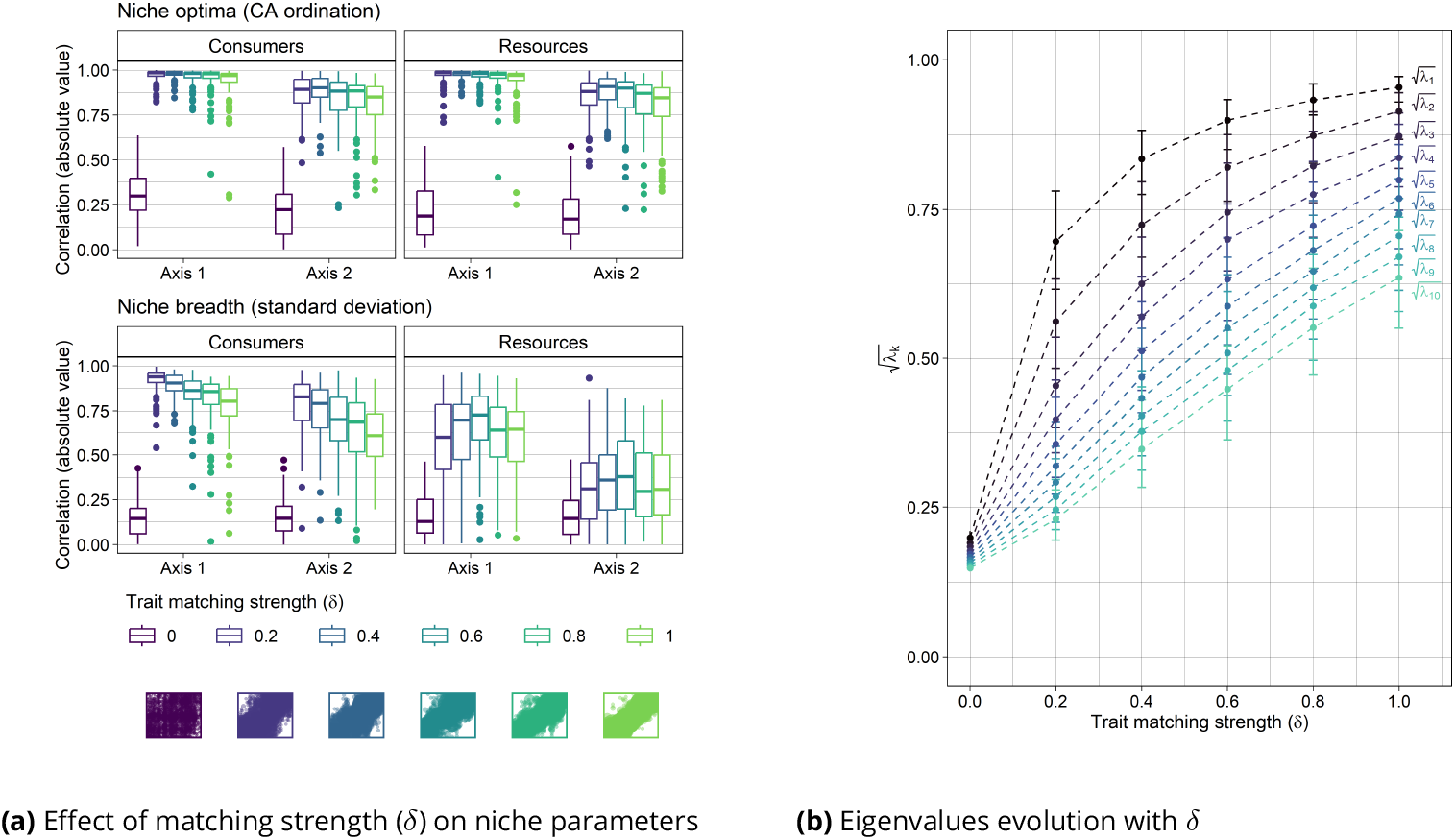
Influence of the strength of trait matching *δ* on niche parameters and latent traits correlations. (a) Quality of the inference of niche parameters depending on values of *δ*. The y-axis is the absolute value of the correlation between true and estimated values of niche optima and niche breadths (respectively top and bottom plots). Matrices exemplify network data (resources in rows, consumers in columns) generated with the corresponding matching strengths. Points represent non-null interactions, with size proportional to the number of observed interactions censored at the 1st and 99th percentiles. (b) Square root of eigenvalues 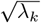 (absolute value of latent traits correlation) depending on different values of *δ*. Colored points and dotted line represent the square root of an eigenvalue, and error bars represent the 2.5 and 97.5-th percentiles on the eigenvalues over 100 repetitions.

The model is robust to skewed abundance distributions, although performance drops slightly with very skewed distributions (Experiment (**n**^**r**^, **n**^**c**^), Figure 3b). Moreover, the performance is better for intermediate niche breadth (Experiment (*µ*_breadth_), Figure 3c). Here, the optimal performance is reached with a niche breadth of 10, representing 10% of the total length of the trait gradient in our setting. Regarding the heterogeneity of consumers’ niche breadths (Experiment (*σ*_breadth_), Figure 3d), homogeneous niche breadths improve the performance. In particular, the niche breadth of resources is correctly recovered only with very homogeneous consumer niches.

Surprisingly, niche parameters are optimally estimated for an intermediate matching strength *δ* (around 0.2) (Figure 4a). However, Figure 4b shows that the square roots of eigenvalues (absolute value of the correlation between latent traits) increase with *δ*. This indicates that a stronger matching signal is inferred as *δ* increases.

### Data analysis

The permutation chi-squared test performed on the interaction table shows that there is a non-random structure in the network (*χ*^2^ = 17784, 2000 permutations, p-value = 5.0 *×* 10^*−*4^). The effect size computed with Cramér’s *V* (*V* = 0.29, *IC*_95_ = [0.26, 1.00]) suggests a small to medium effect (Ellis, 2010).

CA eigenvalues suggest that matching between latent traits is quite strong. Here, for the first three eigenvalues, we have 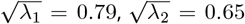 and 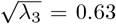, which correspond to the absolute value of the correlation between latent traits (respectively on axes 1, 2 and 3).

With reciprocal scaling, we can position interactions in the multivariate space and use them to visualize species niches. On Figure 5, interactions (points) are grouped by interacting partner. Figures 5a and 5b show species’ niches by grouping interactions by bird and plant species, respectively. The first 2 axes together explain 28.7% of the variance in the data (17.2% on axis 1 and 11.5% on axis 2). Eigenvalues suggest that the third axis also holds structure (10.9% variability), but here we do not discuss this axis for concision.

**Figure 5.**
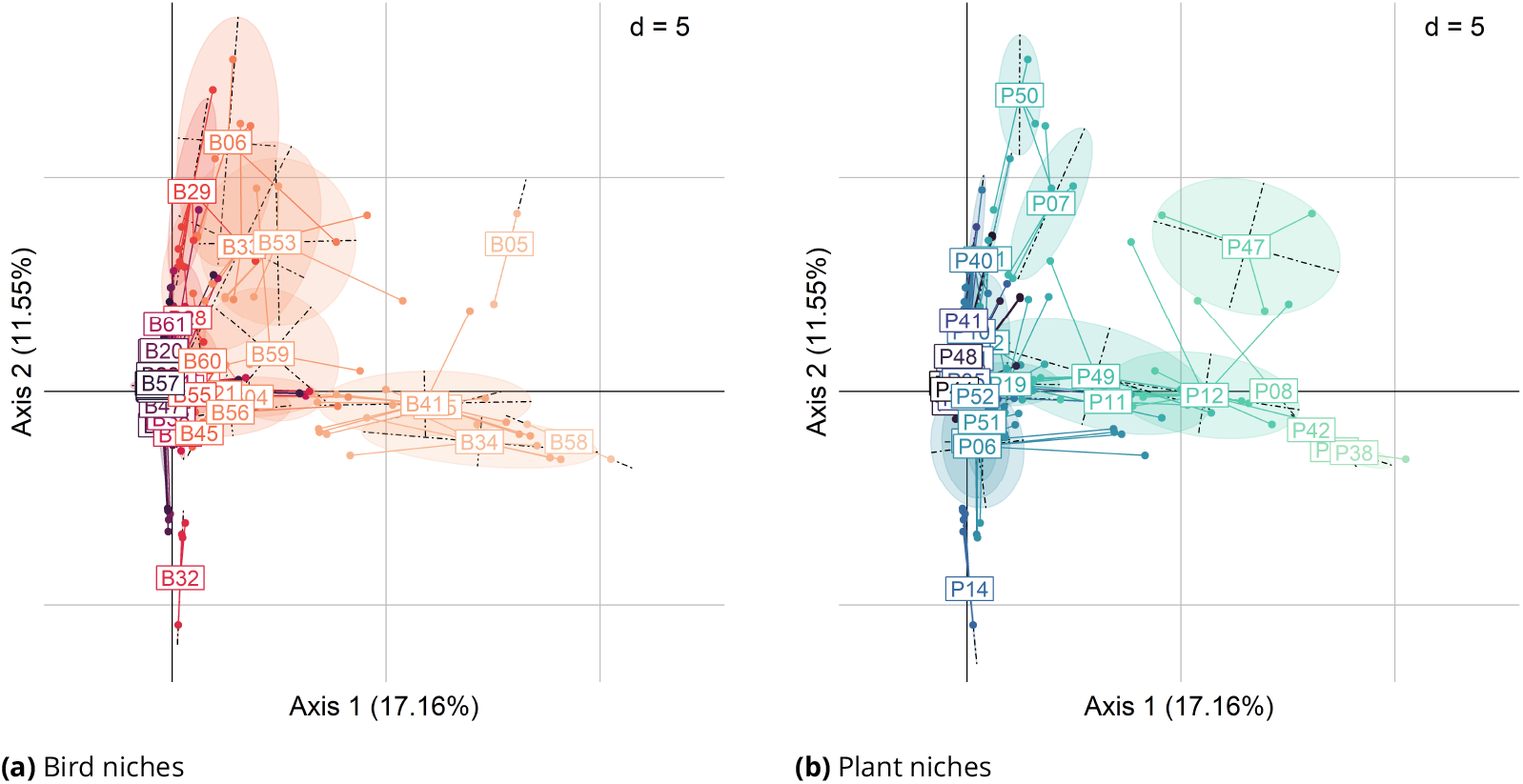
Reciprocal scaling of the birds-plants interaction network. Points correspond to the interaction scores *h*_*k*_(*i, j*) in the first 2 dimensions, grouped by bird (a) or plant species (b). Species label are placed at the reciprocal scaling mean, and ellipses correspond to the bivariate normal distribution with variances and covariances given by reciprocal scaling (with a scaling factor of 1.5, i.e. the ellipse axes lengths are equal to 1.5 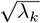 on axis *k*, corresponding to around 67% of the points contained in the ellipse). Ellipses are colored according to the rank of their centroid on the first axis. The correspondence between species’ codes and names is given in Appendix D.

On these figures, two birds (or plants) whose means are located nearby have similar niche optima. Moreover, a bird and a plant that interact preferentially have close niche optima. For instance, the interaction between B05 (*Cephalopterus ornatus*) and P47 (*Symplocos arechea*), which are close to each other, represents respectively 66% and 33% of their total interactions.

These figures also show how niches are distributed in the latent traits space. For birds (Figure 5a), many have a niche located around the origin. These birds interact with average plants, in the sense of their latent traits. Then, we can distinguish two clusters along the positive portions of the two main axes. They show two strategies of birds along the two latent plant traits axes: either interact with plants with above-average values on axis 1, and small values on axis 2, or the reverse. This open triangle is completed by a few bird species around the diagonal (e.g. B33 *Penelope montagnii*, B53 *Thraupis bonariensis* and B59 *Turdus chiguanco*), which interact with plants with average latent traits on axis 1 and above-average latent traits on axis 2.

For plants, many also have a niche optimum close to the origin (they interact with average birds). Like for birds, two clusters emerge along the two main axes, which also reflects two plant strategies. However, contrary to birds, we cannot clearly see plant species around the diagonal, except P47 (*Symplocos arechea*).

To investigate the drivers of niche breadth, we modeled the relationship between species’ niche breadths and their latent traits, also corresponding to their niche optima (Figure 6). On latent trait axis 1, for birds, there is a concave relationship with a positive linear component (*R*^2^ = 0.73), but the variance of the residuals increases along axis 1. For plants, there is only a concave relationship (*R*^2^ = 0.77). On axis 2, for birds, we have a convex relationship (which seems to be driven mainly by B32 *Patagioenas plumbea*) with a strong linear positive component (*R*^2^ = 0.70). For plants, we have a very weak relationship (*R*^2^ = 0.21) with linear positive and weak convex components.

**Figure 6.**
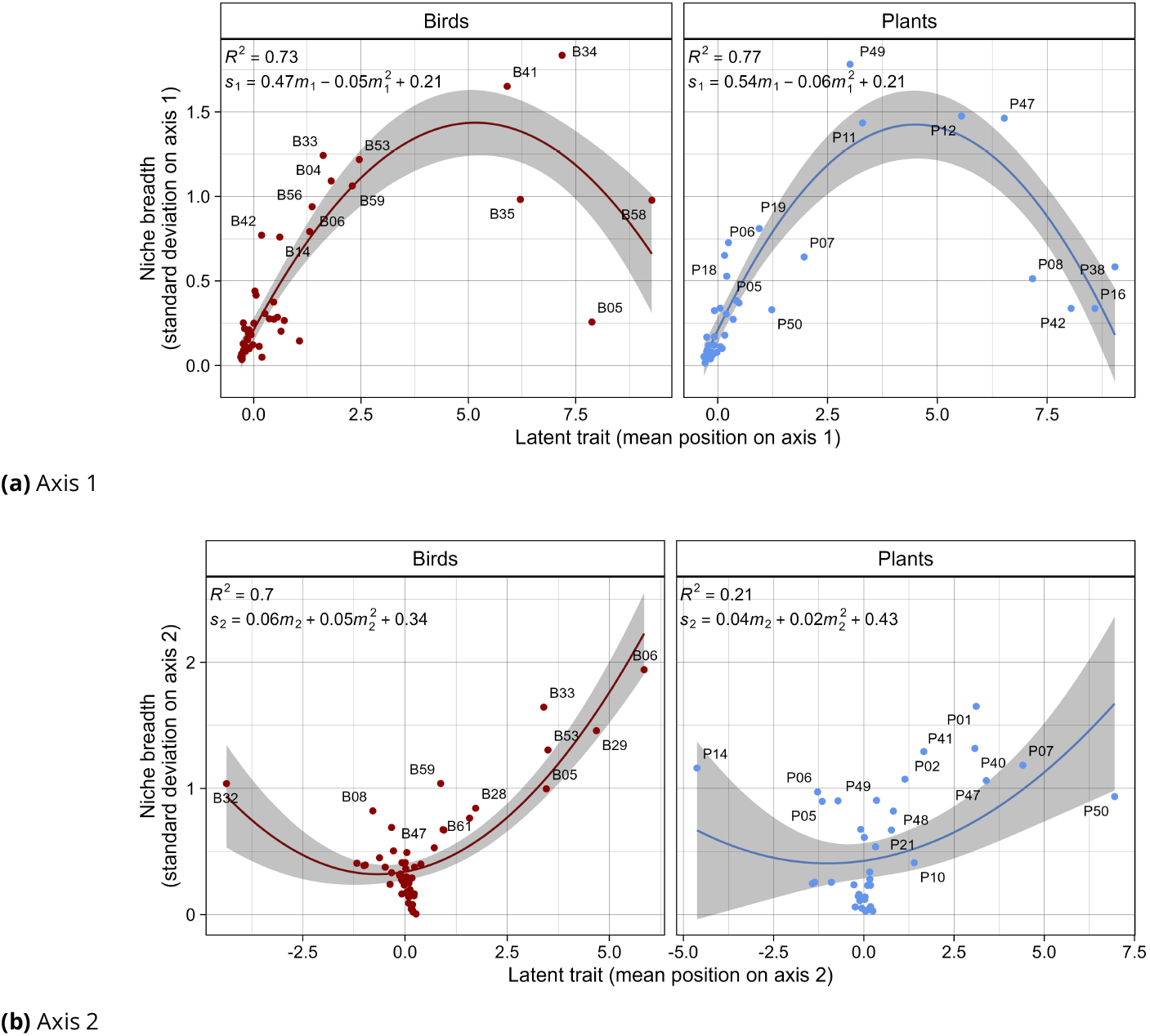
Niche breadth versus niche optimum on the 2 multivariate axes. Niche breadths and optima were computed with reciprocal scaling. The top graphs (a) shows the relationship on axis 1 and the bottom graphs (b) on axis 2. The left and right panels represent bird and plant species, respectively. The solid line is the prediction of the best linear model and the gray shading represents the 95% confidence interval around the predicted mean value. The coefficient of determination (*R*^2^) and the linear model equation are written in the top left corner.

To interpret the latent trait axes, we correlated measured and latent traits a posteriori (Figure 7). All traits correlate positively with axis 1, strongly suggesting that this latent trait represents a size effect. Axis 2 is characterized by its negative correlation with plant height, Kipp’s index and crop mass. All plant traits are also strongly correlated with each other. For birds, body mass and bill width are highly correlated. Regarding cross-trophic level traits, Kipp’s index is strongly positively correlated with crop mass and plant height.

**Figure 7.**
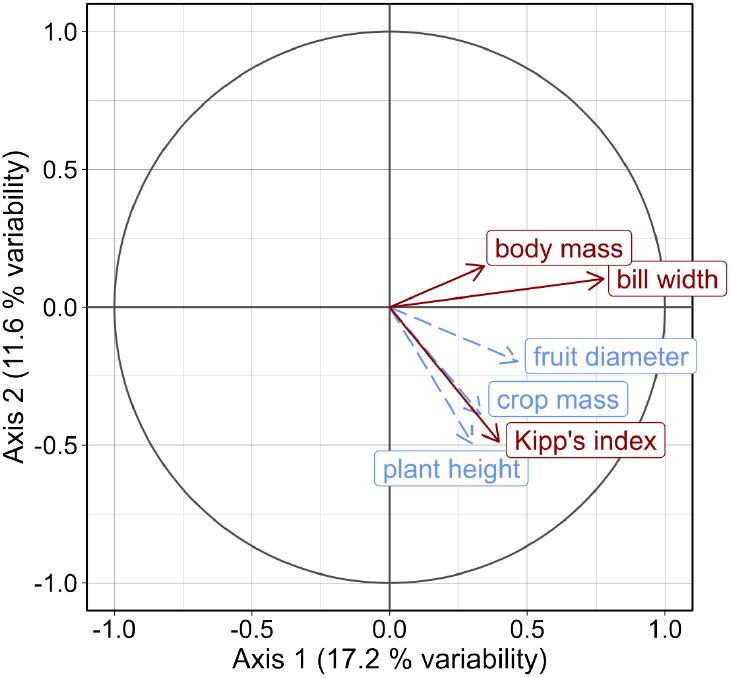
Correlation circle measured a posteriori between species measured and latent traits. Bird traits are shown in solid red lines and plant traits in dashed blue lines. Axis 1 shows a size effect while axis 2 is defined by the matching between plant height and Kipp’s index.

## Discussion

In this article, we used CA and reciprocal scaling on both simulated and real data to quantify species interaction niches. With the case study, we also showed how this approach can be used to test hypotheses on the drivers of species position on the specialist-generalist gradient.

### Simulation

The simulation study shows that the method can recover niche optima, but results are mixed regarding niche breadth. In particular, the niche breadths of resource species are poorly recovered overall, except in a few particularly favorable cases.

Unsurprisingly, performance increases with sampling intensity (Figure 3a). Moreover, when the proportion of variation explained by traits is smaller (on axis 2 of the simulation study), performance drops.

The estimation is robust to skewed abundances, but performance decreases with extremely skewed data (Figure 3b): in that case, we can hypothesize that the niches of rare species are estimated less precisely because they have very few observed interactions.

Consumers’ niche breadth affects the inference of niche parameters (Figure 3c). Indeed, as species are ordered using their similarity in interacting partners, CA needs some overlap in interactions to position them on the latent trait axis. Therefore, there is a trade-off between narrow and wide niches: they need to overlap enough to order adjacent species, but not too much so that there is still a structure in the network.

The heterogeneity of consumers’ niche breadth decreases the performance of the estimation (Figure 3d). We can hypothesize that this is due to insufficient overlap between specialists’ and other species’ niches. This might bias the placement of niche optima for specialists, and the estimation of niche breadths for their interacting partners.

Surprisingly, a stronger trait matching does not always yield a better recovery of niche parameters (Figure 4a). As expected, when there is no trait matching (*δ* = 0), niche parameters are poorly recovered, because the true niche is random. However, the performance is best when the matching strength is intermediate (*δ* = 0.2), and decreases with stronger matching, although it remains quite good even when matching is maximal (*δ* = 1). This can be explained by the requirements of CA to perform latent ordination. Indeed, if the matching is too strong, the niches of species become increasingly separated in the ordination space. Strong trait matching essentially translates into smaller niche overlap. In such cases, the small overlap between species’ interacting partners makes species ordination difficult (as discussed above for the heterogeneity of consumers’ niche breadths). Here, when *δ* = 1, all eigenvalues are also closer to 1 (Figure 4b): the structure of the network is recovered in more than two axes, whereas the true structure used to generate data is driven by two matching traits (i.e. two axes). For this reason, the latent traits on the first two axes are less close to the two traits used in the simulation, thus yielding less accurate estimations of the niche parameters. Therefore, if the CA of an interaction network yields eigenvalues that are all close to one, species’ latent traits should be interpreted with caution. In that case, one can reasonably conclude that there is a strong trait matching, but suspect that the dimensionality of trait matching is lower than the one inferred.

Finally, niche parameters are better recovered for consumers than for resources, especially niche breadths. This might arise from the asymmetric way niche breadths are specified in our simulation model. Indeed, whereas consumers’ niche breaths are specified in the model, niche breadths for resources are driven entirely by the niche breadth of the consumers they interact with.

Our simulation model makes a number of simplifying hypotheses about interaction networks. First, only consumers have a preferred niche breadth. This makes sense in our case study with birds (as consumers) and plants (as resources), because birds’ cognitive processes and movement abilities allow them to choose the plants with which they interact. However, on an evolutionary timescale, plants might evolve to attract more diverse or more similar birds. Specifying a model taking into account both niche breadths requires adapting the trait matching step (Equation (7)), which is difficult. In fact, to our knowledge, no such model exists in the literature. The model also specifies trait matching with a multivariate normal distribution, which can be unrealistic. Despite these limitations, we think that the simulation model used here is useful to provide a first evaluation of the method’s ability to recover niche parameters.

### Data analysis

Regarding real data analysis, a permutation chi-squared test suggests that there is a non-random structure in the network. However, the network is also very noisy, as suggested by visual examination of the network and the small to medium effect size measured with Cramér’s *V*.

This non-random structure is quantified using the chi-squared statistic. However, it uses the matrix margins to assess the expected number of interactions, thus assuming that the number of times a species is observed in the network is a surrogate of its abundance. This hypothesis may not be valid due to sampling effects (Blüthgen, Fründ, et al., 2008; Fründ et al., 2016). In particular, the interactions range of the most abundant species might be better sampled, thus artificially increasing their generality. Reciprocally, under-sampling of rare species might artificially increase their specialization. Moreover, some species engaging in frugivory interactions were probably not sampled, which may bias niche estimates.

In the results, we explored the niche patterns for the first two axes (or latent traits). These two axes represent 28.7% of the total variation in the network, so the patterns discussed below explain only a small part of the network structure.

To interpret latent traits, we correlated them with measured traits. The first latent trait (axis) is mostly correlated with birds and plants size. There is a concave relationship with a positive linear component between birds’ niche breadth and the first latent trait (Figure 6a, left). This suggests that the larger birds are, the wider their niches are, but that medium-sized birds have the widest niches. We can hypothesize that birds are limited by beak size: small-billed birds may not be able to grasp large fruits, while larger-billed birds can swallow even small fruits (Wheelwright, 1985). However, birds that are too large do not eat very small fruits, because they need to satisfy their high absolute energy expenses. Therefore, they focus on fruits that are larger, or more abundant on the plant to minimize foraging time (Albrecht et al., 2018; Schoener, 1971). This limiting effect of energy requirements is weak: our results rather suggest a diversity of strategies for large birds. For instance, B34 (*Pharomachrus antisianus*) and B41 (*Rupicola peruvianus*) eat fruits of diverse sizes. On the contrary, B05 (*Cephalopterus ornatus*) and B58 (*Trogon personatus*) choose fruits of more uniform sizes. In addition, the correlation between birds body mass and plant productivity indices, such as crop mass or fruit diameter, is weak (Figure 7). This may be due to the fact that birds’ foraging is partially modulated by other traits, like whether they are fruit-gulpers or pulp-feeders (Palacio et al., 2017).

For plants (Figure 6a, right), the concave relationship between niche breadth and the first latent trait (axis) is even clearer. This suggests that small plants attract only small birds, plants of intermediate size attract birds of diverse sizes, and large plants attract large birds. Generalist plants have diverse strategies: P11 (*Elaeagia mariae*) and P49 (*Turpinia occidentalis*) predominantly interact with small and medium-sized birds, while P12 (*Endlicheria sp*.) and P47 (*Symplocos arechea*) interact with medium-sized to large birds. A closer examination of their respective traits reveal that the first latent trait of P11 and P49 is predominantly driven by their height, which does not impose a harsh barrier to interactions for small birds. In contrast, P12 and P47 produce fruits larger than the mean bill width, so the interaction barrier is stronger.

On axis 2, we interpret the relationship for birds only Figure 6b, left), because the relationship for plants is very weak. We focus on the linear component between bird’s latent traits and niche breadths, because the convexity seems to be driven by a single species (B32, *Patagioenas plumbea*). The second latent trait is negatively correlated with Kipp’s index (wing pointedness) and plant height. This means that birds with pointed wings consume fruits from plants of homogeneous heights, whereas birds with rounded wings interact with plants of diverse heights. Rounded wings confer more maneuverability in dense vegetation (Thiel et al., 2023), so birds with rounded wings primarily feed on small plants in the dense understory. Conversely, birds with pointed wings preferentially forage on tall plants in the more open canopy. Here, we find a higher specialization for birds with pointed wings: these birds might find it difficult to navigate dense understory, while birds with rounded wings may move to the canopy at a lesser cost (Thiel et al., 2023). Canopy birds might also focus on highly productive fruit patches by choosing plants with high crop yields, as suggested by the high correlation between crop mass and Kipp’s index (Foster, 1990; Thiel et al., 2023). The higher specialization of canopy birds seems to contradict the findings of Schleuning et al. (2011): however, in this study, the authors measured niche breadth as the diversity of interacting partners, not the diversity of their (latent) traits.

A limitation of the results discussed above is that the network analyzed here is an aggregation of four observed samples collected over a year. Therefore, the inferred niche is a mean of the interactions a species engages in throughout the year, although it may vary dynamically depending on various temporal processes (Poisot et al., 2015; Singer and McBride, 2012). Moreover, the two axes (latent traits) discussed above explain roughly 30% of the variability, so they account for a small fraction of the non-random network structure. Discussing patterns on the other axes would be important, in particular on the third axis which explains almost the same variability as the second axis. Eigenvalues also suggest that underlying structures are found up to the 6th axis, and considering 6 latent traits would explain more than 60% of the variation of the network (see Appendix E).

### Conclusion and perspectives

This article describes correspondence analysis and reciprocal scaling applied to ecological networks. These methods are well-suited for exploratory analyses, and not limited by costly trait collection. CA allows to evaluate a potential signal and the dimensionality of trait matching (Eklöf et al., 2013), without requiring trait data. Reciprocal scaling provides a quantitative measure of the interaction niche as a hypervolume, using a Gaussian approximation. The resulting niche is a hyperellipsoid in *n* dimensions, corresponding to the isocontours of a multivariate normal distribution.

Our framework links trait matching and species specialization-generalization, through the notion of interaction niches (Albrecht et al., 2018; Dehling, Lai, et al., 2025; Godoy et al., 2018). Trait matching is viewed as the alignment between species niche optima, defined by their (latent) traits. Specialization-generalization are viewed as the diversity of (latent) traits of a species’ interacting partners.

CA and reciprocal scaling are proposed as a powerful tool for analyzing species interactions, but they are also limited within a specific scope. First, latent traits conflate several processes, like species’ spatial and temporal variations in abundances, phylogeny, or sampling biases. Therefore, the estimated niche corresponds to species’ realized niche, i.e. the interacting partners a species effectively interacts with (Devictor et al., 2010; Hutchinson, 1957), which can be influenced by a variety of factors beyond their traits. For instance, the distribution of traits in the community might constrain species to settle for suboptimal interacting partners. Additionally, competition may also constrain species to partition their realized interaction niches. In Appendix A, we show that the method can also recover fundamental niches (although not as well as realized niches). Second, reciprocal scaling estimates a Gaussian approximation of the species niche. While this approximation will likely be sensible for symmetric and unimodal niches, the Gaussian estimate might be irrelevant for species with highly skewed or multimodal niches (Benadi et al., 2022; Blonder et al., 2014). For example, Gaussian niches would likely be a poor estimate of asymmetric niches constrained by exploitation barriers (Santamaría and Rodríguez-Gironés, 2007).

The dimensionality of latent traits estimated with CA depends on the dimensionality of the original data: it is min(*r* − 1, *c* − 1). Therefore, when interpreting data, it is necessary to choose the number latent traits to retain. In practice, ecologists should explore the first *k* axes (latent traits), until eigenvalues associated to these traits decrease abruptly (i.e. the gain in explained variation becomes negligible). They may also use methods guiding the choice of dimensionality in multivariate analyses (similar to the method proposed by Dray, 2008, for principal component analysis).

Niche breadths have been studied using other measures in the literature, often based on the identity of interacting partners, as opposed to their traits (e.g. the index *d*^*′*^, which also takes into account species availability, proposed by Blüthgen, Menzel, et al., 2006). A notable exception is the process-related niche proposed by Dehling and Stouffer (2018), which measures the functional diversity of a species’ interacting partners using their traits projected in a multivariate space. Reciprocal scaling bears similarities with both methods, as it quantifies niche breadth from the interaction matrix only, but also uses latent traits. Our measure hence provides a complementary way to define niche breadth, that takes into account both species characteristics (using latent traits) and their availability (using species’ relative frequencies in the niche breadth measure).

CA and reciprocal scaling could be extended in various ways. First, we could measure niche overlap by quantifying the ellipses overlapping area (similarly to Pappas and Stoermer, 1997). This would allow to gain insights regarding the similarity, or dissimilarity, of species niches, thereby potentially allowing to measure competition strength. Second, it would be interesting to estimate a confidence interval around niche breadth and optima, for instance using a bootstrapping approach (i.e. iteratively removing species from the network and estimating niche parameters on these bootstrapped data).

Although the latent approach of CA has many benefits, when traits are known, it would also be interesting to constrain the multivariate axes with traits. Such constrained analyses have been performed with RLQ or fourth-corner analyses in the context of network data (Albrecht et al., 2018; Bender et al., 2018; Dehling, Töpfer, et al., 2014). However, fourth-corner analysis considers only pairs of traits, and does not ordinate species in the multivariate space. On the other hand, RLQ is covariance-based, which does not allow to partition the variation explained by traits in interaction networks.

An alternative would be to use constrained versions of CA. More precisely, we could use canonical correspondence analysis (ter Braak, 1986) and double constrained correspondence analysis (ter Braak et al., 2018). These canonical methods would allow to quantifying the part of variation explained by trait matching in ecological networks. They would also eliminate the need for important niche overlap, which is necessary to recover structuring trait axes with latent gradients in CA. To our knowledge, constrained versions of correspondence analyses have never been applied to interaction networks. Moreover, for all methods above, there is no known way to measure niche breadth. The part of variation explained by trait matching would also likely be biased, as this is the case for canonical correspondence analysis (Peres-Neto et al., 2006) and fourth-corner analysis (Dray and Legendre, 2008). Therefore, further methodological developments in multivariate analyses could benefit network analysis.

## Appendices

### A Realized and fundamental niches

In the article, we measured reciprocal scaling performance by comparing the inferred niche parameters to the realized niches. However, we could also compare the inferred niche parameters to the fundamental niches, i.e. the trait values (and niche breadths, for consumers) fixed in the model. Below, we present the performance of the model when we consider the fundamental niches as the ground truth (results are shown only for consumers). For simulation parameters, we choose the same parameters as described in Table 1.

**Figure A.1.**
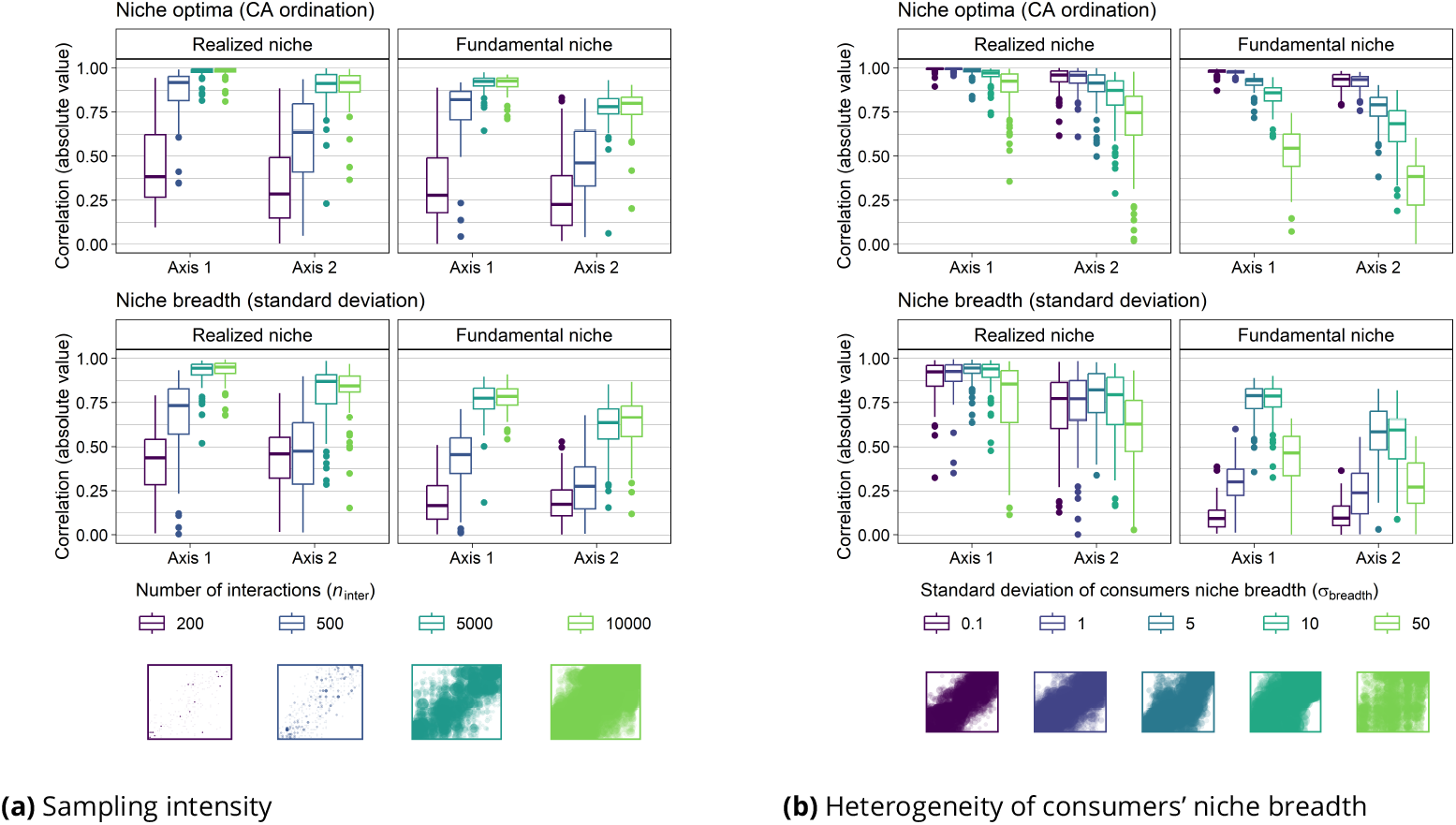
Performance of the model to infer fundamental niche parameters for consumers. (a) Shows the effect of the sampling intensity and (b) shows the effect of the heterogeneity of consumers’ niche breadths on the model performance. The y-axis is the absolute value of the correlation between true and estimated values of niche optima and niche breadths (respectively top and bottom of each subplot). The correlation is presented separately for realized and fundamental niches (respectively left and right panels) on axes 1 and 2. Below the legend, matrices exemplify network data (resources in rows, consumers in columns) generated with the corresponding parameter value. Point sizes are explained in Figure 3.

In most cases, the performance of the model to recover fundamental niche is a reflection of the realized niche performance, but with slightly worse performance. As an example, we present the model performance for realized and fundamental niches depending on sampling intensity in Figure A.1a.

However, the performance pattern is different when the true niche breadth varies: whereas the realized niche is less well recovered as the standard deviation of niche breadth increases, the fundamental niche is better recovered for intermediate values of niche breadths (Figure A.1b). We can see it as an artifact of the performance metric used here: when niches breadths are very homogeneous, the model is able to recover the absolute niche breadths, but not their relative order because they are very homogeneous.

### B Networks visualization using CA

Correspondence analysis can also be useful to reorder rows and columns of the analyzed interaction matrix. Many ways to visualize interaction networks have been proposed, with all of them highlighting a specific aspect of network structure like centrality, modularity or nestedness (Araujo et al., 2010; Miele et al., 2019). CA can also serve this purpose. The figure below shows the interaction matrix corresponding to the bipartite birds-plants interaction network analyzed in this article. In Figure B.1a, rows and columns are ordered according to the alphabetical order, while in Figure B.1b they are ordered according to their rank on the first axis of the CA. This means that species are positioned in an ecologically meaningful way, according to their interaction niche optima on the first latent trait axis. Here, we can see two weakly connected modules, corresponding to the bulk of small plant and bird species that interact together (bottom-left corner), and fewer large birds and plants that also interact together (top-right corner). A few species interact with species from both modules (like P12 (*Endlicheria sp*.) and B41 (*Rupicola peruvianus*)), and correspond to the generalist intermediate species from Figure 6.

**Figure B.1.**
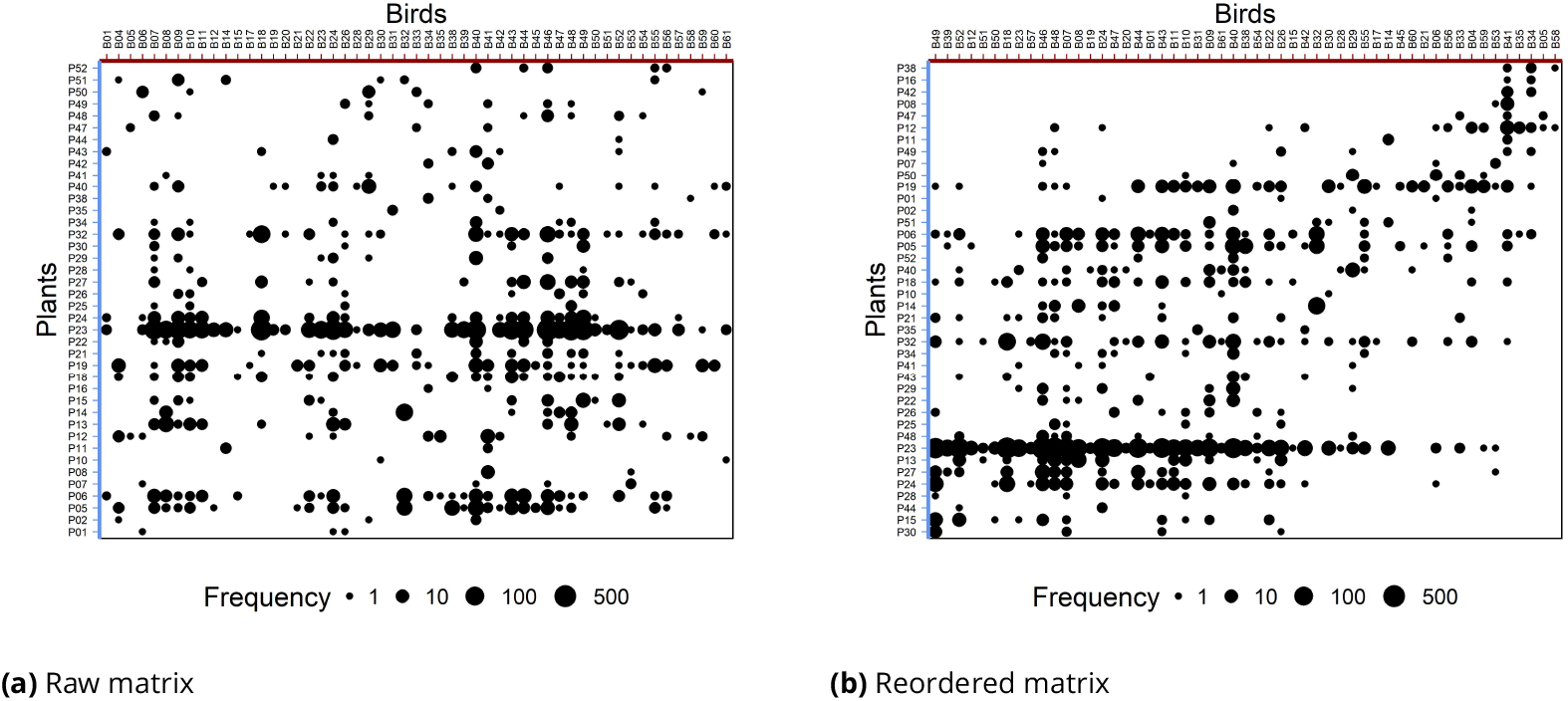
Interaction matrix reordered using correspondence analysis. (a) Rows and columns are ordered alphabetically. (b) Rows and columns are ordered following the coordinates of species on the first axis of CA. Points represent non-null interactions, with size proportional to the number of observed interactions on a log scale.

### C Reciprocal scaling formulas

Reciprocal scaling allows to compute niche optima and niche breadths on each axis *k*, as well as the covariance between two niche axes *k* and *l*. Each of these quantities can be expressed either with the scores **h**_*k*_ derived from reciprocal scaling or equivalently using the scores 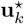 and 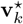 of correspondence analysis, so each quantity has two equivalent expressions. The corresponding formulas are written below, adapted from Thioulouse and Chessel (1992) to match our notation.

#### Niche optima

Niche optima can be expressed either as a weighted mean of the correspondences scores, or from the CA scores. For the resource species *i*, we have:

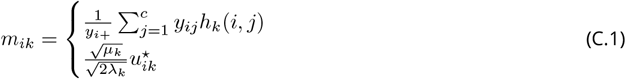

where 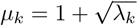.

Similarly, for consumer species *j*, we have:

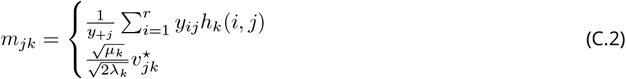

#### Niche breadths

The niche breadth can be expressed as the weighted standard deviation of the correspondences scores or from the CA scores as well. For resource species *i*, we have:

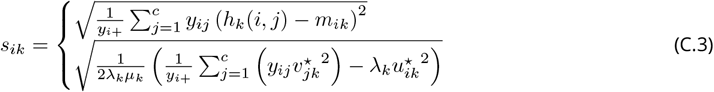

Similarly, for consumer species *j*, we have:

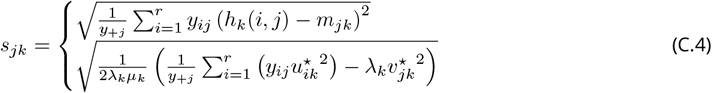

#### Covariances

We can express the covariance between two niche axes from the reciprocal scaling of the CA scores as well. The covariance between niche axes *k* and *l* for resource species *i* is written:

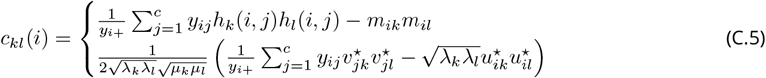

For consumer species *j*, the covariance is written as:

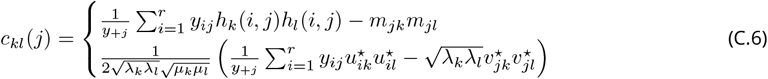

### D Species names

Throughout the article, we use codes to describe species. Below is the correspondence between these codes and the species names, alongside with their traits (all data come from the ANDEAN frugivory dataset (Dehling, Bender, et al., 2021), network Peru1).

**Table D.1.**
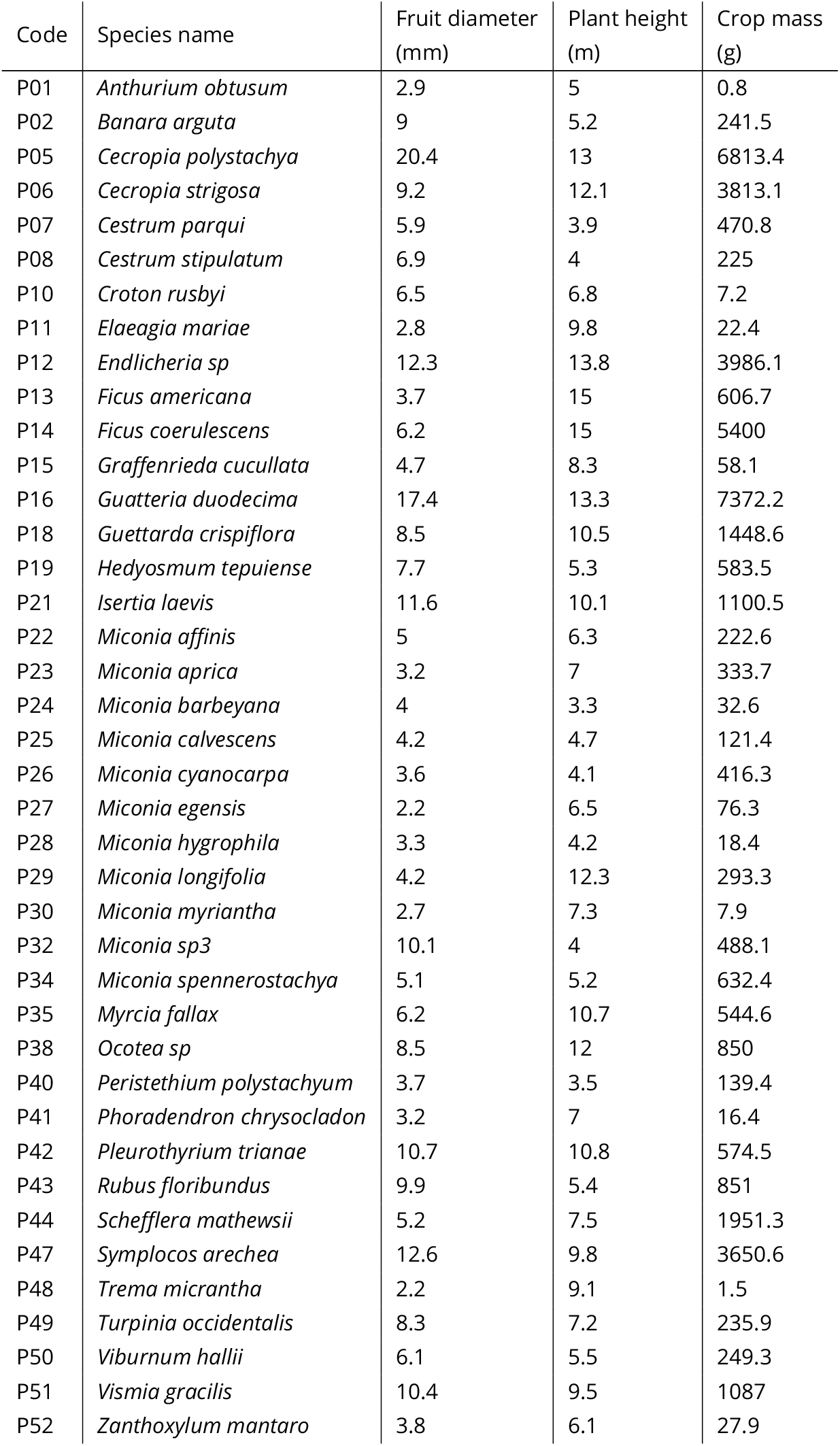
Correspondence between plant codes and species names.

| Code | Species name | Fruit diameter (mm) | Plant height (m) | Crop mass (g) |
| --- | --- | --- | --- | --- |
| P01 | *Anthurium obtusum* | 2.9 | 5 | 0.8 |
| P02 | *Banara arguta* | 9 | 5.2 | 241.5 |
| P05 | *Cecropia polystachya* | 20.4 | 13 | 6813.4 |
| P06 | *Cecropia strigosa* | 9.2 | 12.1 | 3813.1 |
| P07 | *Cestrum parqui* | 5.9 | 3.9 | 470.8 |
| P08 | *Cestrum stipulatum* | 6.9 | 4 | 225 |
| P10 | *Croton rusbyi* | 6.5 | 6.8 | 7.2 |
| P11 | *Elaeagia mariae* | 2.8 | 9.8 | 22.4 |
| P12 | *Endlicheria sp* | 12.3 | 13.8 | 3986.1 |
| P13 | *Ficus americana* | 3.7 | 15 | 606.7 |
| P14 | *Ficus coerulescens* | 6.2 | 15 | 5400 |
| P15 | *Graffenrieda cucullata* | 4.7 | 8.3 | 58.1 |
| P16 | *Guatteria duodecima* | 17.4 | 13.3 | 7372.2 |
| P18 | *Guettarda crispiflora* | 8.5 | 10.5 | 1448.6 |
| P19 | *Hedyosmum tepuiense* | 7.7 | 5.3 | 583.5 |
| P21 | *Isertia laevis* | 11.6 | 10.1 | 1100.5 |
| P22 | *Miconia affinis* | 5 | 6.3 | 222.6 |
| P23 | *Miconia aprica* | 3.2 | 7 | 333.7 |
| P24 | *Miconia barbeyana* | 4 | 3.3 | 32.6 |
| P25 | *Miconia calvescens* | 4.2 | 4.7 | 121.4 |
| P26 | *Miconia cyanocarpa* | 3.6 | 4.1 | 416.3 |
| P27 | *Miconia egensis* | 2.2 | 6.5 | 76.3 |
| P28 | *Miconia hygrophila* | 3.3 | 4.2 | 18.4 |
| P29 | *Miconia longifolia* | 4.2 | 12.3 | 293.3 |
| P30 | *Miconia myriantha* | 2.7 | 7.3 | 7.9 |
| P32 | *Miconia sp3* | 10.1 | 4 | 488.1 |
| P34 | *Miconia spennerostachya* | 5.1 | 5.2 | 632.4 |
| P35 | *Myrcia fallax* | 6.2 | 10.7 | 544.6 |
| P38 | *Ocotea sp* | 8.5 | 12 | 850 |
| P40 | *Peristethium polystachyum* | 3.7 | 3.5 | 139.4 |
| P41 | *Phoradendron chrysocladon* | 3.2 | 7 | 16.4 |
| P42 | *Pleurothyrium trianae* | 10.7 | 10.8 | 574.5 |
| P43 | *Rubus floribundus* | 9.9 | 5.4 | 851 |
| P44 | *Schefflera mathewsii* | 5.2 | 7.5 | 1951.3 |
| P47 | *Symplocos arechea* | 12.6 | 9.8 | 3650.6 |
| P48 | *Trema micrantha* | 2.2 | 9.1 | 1.5 |
| P49 | *Turpinia occidentalis* | 8.3 | 7.2 | 235.9 |
| P50 | *Viburnum hallii* | 6.1 | 5.5 | 249.3 |
| P51 | *Vismia gracilis* | 10.4 | 9.5 | 1087 |
| P52 | *Zanthoxylum mantaro* | 3.8 | 6.1 | 27.9 |

**Table D.2.**
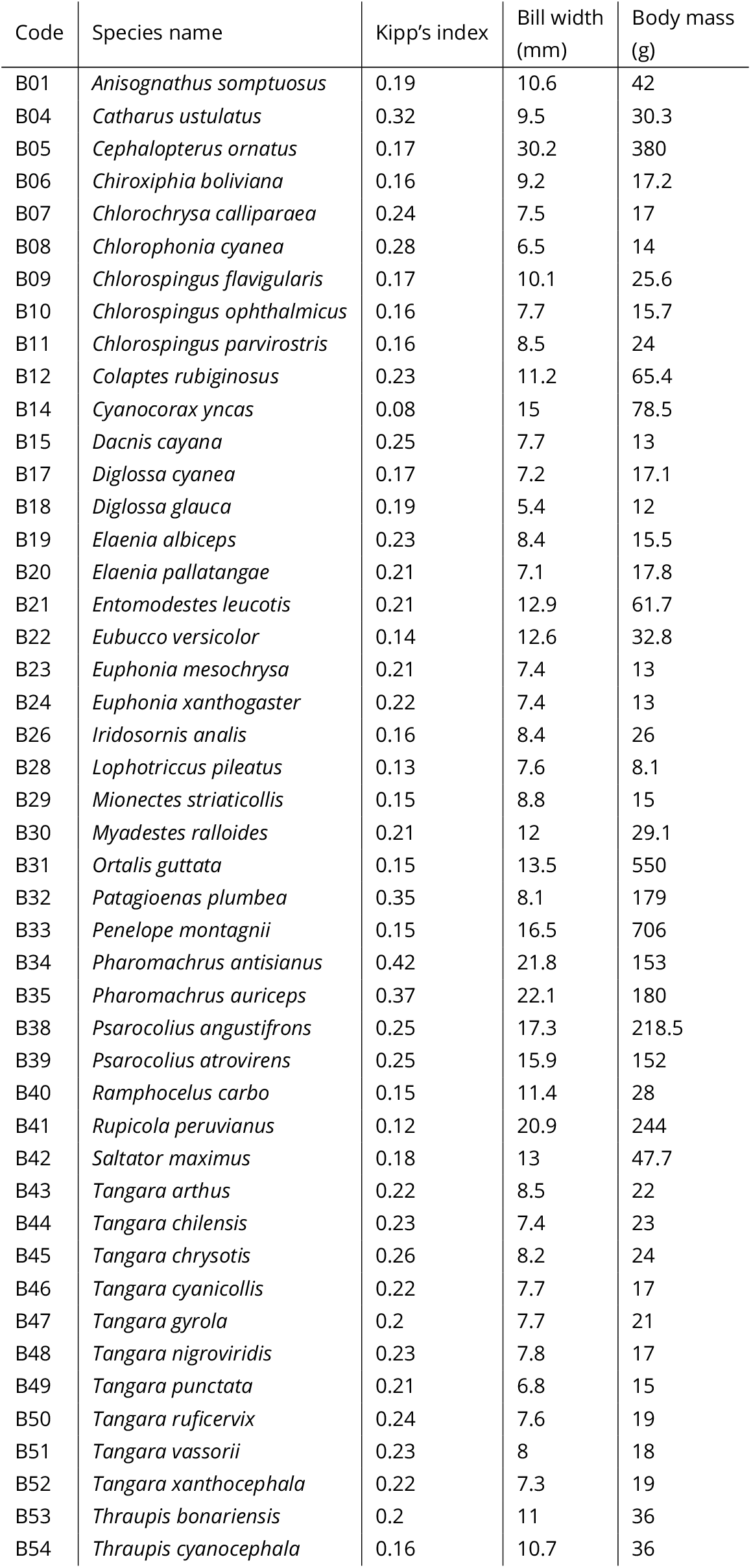
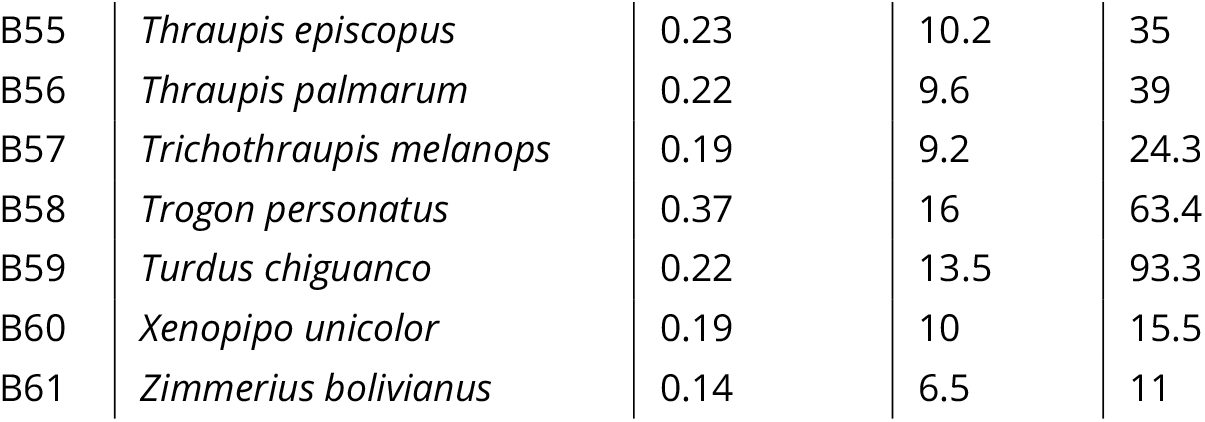
Correspondence between bird codes and species names.

### E Eigenvalues

The Figure below shows quantities related to the eigenvalues of the CA on the example dataset analyzed in the main text (Dehling, Bender, et al., 2021).

Figure E.1a shows the square root of eigenvalues, which represent the absolute correlation between latent traits. We can see a gap between the 3rd and 4th and the 6th and 7th eigenvalues, thus suggesting that underlying structures are found up to the 6th axis.

Figure E.1b shows eigenvalues relative cumulative sum, which represents the part of variation explained by eigenvalues 1 to *k*. Considering the first 6 axes (i.e. 6 latent traits) would explains more than 60% of the variation of the network.

**Figure E.1.**
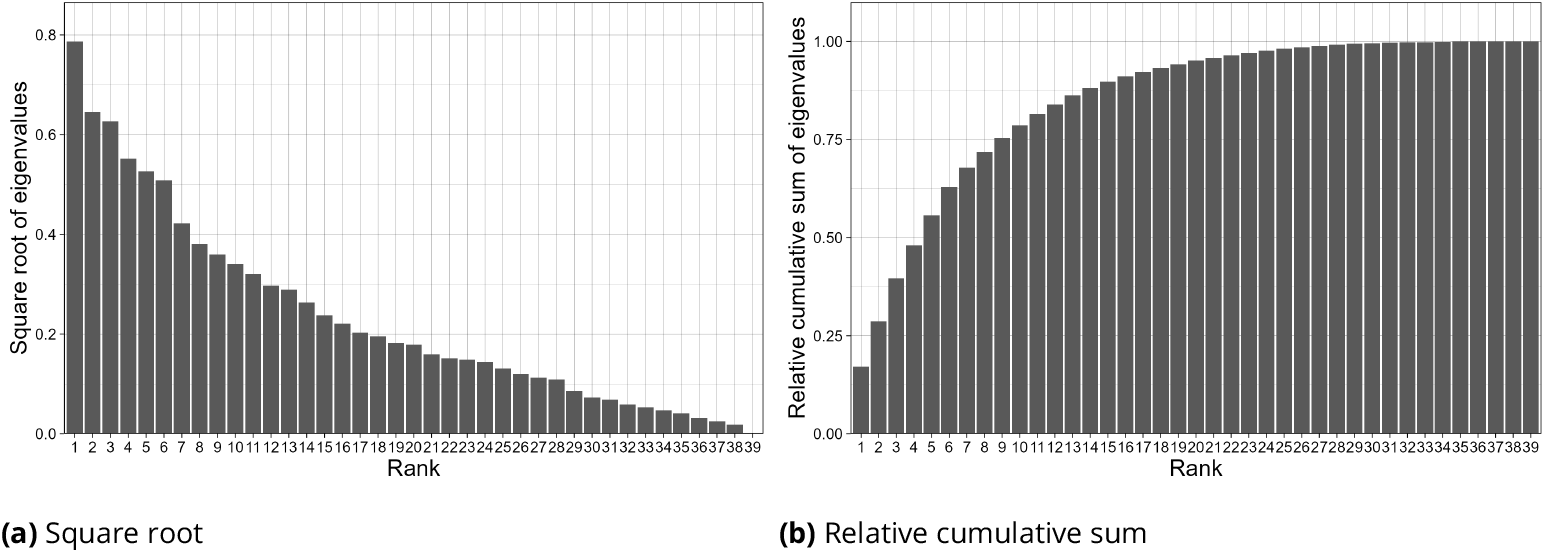
Correspondence analysis eigenvalues. (a) Square root of eigenvalues, which represents the absolute value of the correlation between latent traits. (b) Relative cumulative sum representing the part of variation explained by eigenvalues 1 to *k*.

## Acknowledgements

We thank Mahendra Mariadassou, Vincent Miele, Sara Puijalon, Stéphane Robin and Jan Venter for their input on this work during PhD committee meetings. We also thank Elisa Thébault, Marie-Pierre Etienne, Simon Chamaillé-Jammes and in particular Anne-Béatrice Dufour for their input on this article initially written as a thesis chapter. We are also grateful to the two reviewers, Pedro Henrique Pereira Braga and Ignasi Bartomeus, for their extremely relevant and detailed comments on the manuscript. Preprint version 3 of this article has been peer-reviewed and recommended by Peer Community In Ecology (https://doi.org/10.24072/pci.ecology.100765; Poisot, 2025).

## Data, script, code, and supplementary information availability

Code and data are available at https://doi.org/10.6084/m9.figshare.28625270.v3 (Nicvert et al., 2025).

